# Synbiotic Yogurt with *Lactobacillus plantarum* and *Stevia rebaudiana*: Physicochemical, Microbiological, and Functional Evaluation

**DOI:** 10.64898/2026.04.08.716962

**Authors:** Prabhat Dhakal, Pooja Chaudhary, Tika Bahadur Karki, Suman Lama

## Abstract

Functional dairy products are increasingly recognized for their ability to provide both essential nutrition and additional health benefits. This study aimed to develop and evaluate a synbiotic yogurt enriched with *Lactobacillus plantarum* as a probiotic and *Stevia rebaudiana* extract (1% w/v) as a prebiotic source. Thirteen lactic acid bacteria (LAB) strains were isolated from fermented dairy and vegetable samples and evaluated for probiotic potential through tests for acid and bile tolerance, hydrophobicity, aggregation abilities, and pathogen co-aggregation. Isolate PG1 (*Lactobacillus plantarum*) demonstrated the highest prebiotic growth stimulation index (49%) in the presence of stevia extract and was selected for yogurt formulation. Yogurt samples were prepared and stored at 4°C for 10 days. Physicochemical properties (pH, titratable acidity, and protein content), microbiological viability, total phenolic and flavonoid content, antioxidant activity (DPPH assay), and sensory attributes were monitored. The synbiotic yogurt (St-Y) showed enhanced functional properties, with a total phenolic content of 16.67 µg GAE/g, a flavonoid content of 6.28 µg QE/g, and 57.84% antioxidant activity. Additionally, it showed improved protein content and superior sensory scores compared to control samples. These findings suggest that S. rebaudiana fortified probiotic yogurt can serve as a nutritious, antioxidant-rich, and sensory-acceptable functional dairy product.

## INTRODUCTION

Advances in the food industry and increasing consumer awareness of healthy and nutritious foods have accentuated the demand for functional foods that provide bioactive and therapeutic properties beyond their nutritional value (Shori et al., 2018). Functional foods are now recognized for their potential to support health and reduce the risk of disease, making them an essential component of modern diets (Jackson et al., 2019; Kaur et al., 2022; Orel & Trop, 2014). Among these, probiotic-enriched foods have gained prominence due to their ability to modulate gut microbiota, thereby improving gastrointestinal and systemic health. Probiotics are defined as “live microorganisms which when administered in adequate amounts confer a health benefit on the host” (Hill et al., 2014). Probiotic organisms require effective delivery vehicles to ensure their viability and functionality upon reaching the gut. Fermented dairy products like yogurt serve as ideal carriers of probiotics and functional ingredients into the digestive system (El-Sayed et al., 2022; Ibeagha-Awemu et al., 2009; Ye et al., 2022). However, ensuring the viability of probiotics (typically 8–9 log CFU g−1) throughout product shelf life and gastrointestinal transit remains a critical challenge (Ahmed et al., 2023; Rodrigues et al., 2019).

To enhance probiotic efficacy, prebiotics (indigestible biopolymers that enhance the activity and viability of beneficial microbiota) are frequently combined with probiotics (Gibson et al., 2017). *Stevia rebaudiana*, a natural sweetener rich in bioactive compounds such as stevioside, rebaudioside A, polyphenols, and flavonoids, which possess antimicrobial, antiviral, and antioxidant properties, is a promising prebiotic candidate. Beyond its sweetness, stevia exhibits antimicrobial, antioxidant, and prebiotic properties, potentially enhancing probiotic survival and metabolic activity (Kunová et al., 2014; Rashwan et al., 2023; Wölwer-Rieck, 2012). The synergy between probiotics and prebiotics in synbiotic formulations can amplify their individual benefits, improving microbial viability during storage and gut colonization (Charalampopoulos & Rastall, 2012; Fayed et al., 2019; Fisberg & Machado, 2015).

Despite the potential of stevia as a prebiotic, its application in synbiotic yogurt remains underexplored, particularly regarding its effects on probiotic growth, product stability, and sensory properties. This study aimed to develop a novel synbiotic yogurt by combining probiotic lactic acid bacteria (LAB) with stevia extract. Specific objectives include (1) isolating and screening probiotic LAB from fermented foods, (2) preparing and characterizing stevia extract, (3) evaluating the growth response of selected LAB to stevia, (4) formulating synbiotic yogurt with optimal stevia concentration, and (5) assessing the product’s physicochemical, microbiological, and functional (phenolic/flavonoid content) and sensory attributes. The findings will contribute to the growing body of knowledge on synbiotic foods, offering a functional dairy product with enhanced health benefits and consumer appeal.

## MATERIALS AND METHODS

### 2.1 Collection of samples

The standardized milk (3% fat and 8% SNF) and pasteurized milk were obtained from the Dairy Development Corporation (DDC). Dried leaf powder of Stevia rebaudiana was collected from the local herbal shop, marketed by Himchuli Foods Packaging Industry. The 16S RNA gene sequenced Lactobacillus plantarum (PG1) was provided by the Microbiology Laboratory of the National College. Tama, Gundruk, Sinki, and Dahi were collected from Kathmandu, Chitwan, and Syangja districts for the isolation of lactic acid bacteria.

#### Isolation of lactic acid bacteria

LABs were isolated using the pour plate method on glucose yeast extract peptone agar medium (GYP) supplemented with 1% calcium carbonate. The fermented food samples were serially diluted up to 10^−6^. One ml of the sample from each dilution was inoculated into a sterile petri plate, and then GYP medium was poured carefully. The plates were then incubated at 37℃ for 48 hours. The isolated colonies were preliminarily selected based on a clear zone around the colonies.

#### Identification of the lactic acid bacteria

For preliminary identification of LAB, cultural, microscopic, and sugar fermentation examinations were performed. The isolated colonies were preliminarily examined based on their size, color, shape, consistency, elevation, and margin. For microscopic examination, Gram’s staining was performed.

#### Screening probiotic potential of LAB isolates

##### 2.1.1 Acid tolerance test

A loopful of LAB inoculum was inoculated in 5 ml of MRS broth and incubated at 37ºc for 24 hours. A loopful of overnight culture was inoculated in De Man–Rogosa–Sharpe (MRS) broth having different pH levels (3.0 and 7.4). All the tubes were incubated at 37ºc for 2 hours and then cultured onto MRS agar plates. The plates were incubated at 37ºc for 48 hours. Acid tolerance was assessed in terms of viable colony counts after incubation (Kaushik et al., 2009; Khalil et al., 2018). Results with higher CFU as compared to the initial were termed as acid-tolerant.

##### 2.1.2 Bile salt tolerance test

A loopful of acid-tolerant LAB inoculum was inoculated in 5 ml of GYP broth and incubated at 37ºc for 24 hours. 200µl of the overnight culture of culture was inoculated in 5 ml of GYP broth supplemented with bile salt at different concentrations (0.15%, 0.3%, and 0.5%). All the tubes were then incubated at 37ºc °C for 24 hours, and the growth (optical density) was measured at OD_600_ with a UV spectrophotometer (Safi et al., 2023). Increased growth (OD) in the presence of 0.3% bile salt, was considered as Bile salt-tolerant isolates.

##### 2.1.3 Phenol tolerance test

A loopful of bile salt-tolerant LAB inoculum was inoculated in 5ml GYP broth and incubated at 37ºc for 24 hours. 200µl of an overnight culture of culture was inoculated in 5 ml of GYP broth supplemented with phenol at different concentrations (0.2% and 0.4%). All the tubes were then incubated at 37ºc °C for 24 hours, and the growth (optical density) was measured at OD600 with a UV spectrophotometer (Safi et al., 2023). Isolates with higher OD values in the presence of 0.4% phenol were considered phenol-tolerant isolates.

### 2.2 Sugar fermentation patterns of probiotic isolates

They also underwent sugar fermentation test in 5 ml fermentation broth supplemented with 1% sugars (Glucose, Lactose, Fructose, Mannose, Sucrose, Maltose, Xylose, Ribose, Rhamnose, Galactose, Starch, and Mannitol), 0.01% indicator phenol red (Breed et al., 1957). The isolates were inoculated separately in fermentative broth, and these tubes were incubated at 37℃ for 36-48 hours. The changes in the color of the media due to the growth and utilization of that sugar by bacteria were observed.

### 2.3 Preparation of an aqueous extract of prebiotics

*Stevia rebaudiana* was first homogenized with distilled water in a 1:10 (w/v) ratio. The solution was then heated at 80ºc for 30 minutes on a hot plate with a magnetic stirrer and allowed to cool down at room temperature. The mixture was filtered using Whatman filter paper (pore size = 11μm). The collected filtrate was finally stored at 4ºc until further use.

### 2.4 Growth stimulation effect of prebiotics

The effect of prebiotic aqueous solution on the growth of probiotics was analyzed using the method described by Sakr & Massoud (2021) with slight modifications. Probiotic isolates were first streaked onto a GYP plate and incubated at 37 °C for 24 hours. The single isolated colony was then inoculated into GYP broth and incubated at 37ºc for 24 hours. The optical density of the probiotic isolate was adjusted to 1.5×10^8^ CFU/ml. Afterward, 1% culture was inoculated into GYP broth and modified GYP broth in which glucose was replaced by either 0.5% or 1%, or 2% of prebiotic extract solution, and incubated at 37º c for 48 hrs. and absorbance reading was taken at 600nm using a spectrophotometer. The growth index was calculated using the formula:

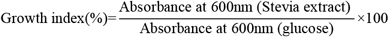

### 2.5 Determination of probiotic properties

#### 2.5.1 Antagonistic activity against pathogenic bacteria

The antimicrobial activity of PG1 (*L. plantarum*) against six different food pathogenic bacteria, such as *Salmonella typhi, S. paratyphi, K. oxytoca, Klebsiella pneumoniae, Pseudomonas aeruginosa*, and *Shigella boydii*, was assessed using an agar well disc diffusion assay. On the MRS plate, *L. plantarum* was initially streaked and then incubated for 48 hours at 37°C. After being inoculated into 5 mL of MRS broth supplemented with 250 mM glycerol, the single isolated colony was cultured for 48 hours at 37°C. After that, the culture was put into an Eppendorf tube and centrifuged for 20 minutes at 4°C at 12000 rpm. Cell-free supernatants (CFS) were kept at 4°C.

On a nutrient agar plate, each food-borne pathogenic bacterium was streaked, and the plate was then incubated at 37°C for 24 hours. *L. plantarum* was separated into a single isolated colony and then inoculated into 5 ml of nutrient broth and cultured for 24 hours at 37°C. 5 ml of actively grown culture was swabbed into a Plate Count Agar (PCA) plate with 6 mm diameter wells. 100 μl of CFS and water (control) was placed into each well and incubated at 37℃ for 24 hrs. A distinct zone of inhibition was observed surrounding the well.

#### 2.5.2 Cell hydrophobicity

##### 2.5.2.1 Bacterial hydrophobicity

The cell surface hydrophobicity to xylene, chloroform, ethyl acetate, hexane, diethyl ether, toluene, and benzene was determined by the method described by Abbasiliasi et al. (2017) with some modifications. *L. plantarum* was first streaked on an MRS plate and incubated at 37℃ for 48 hours. The single isolated colony was inoculated into 5 ml of MRS broth and incubated at 37℃ for 48 hours. The culture was then centrifuged at 8,000 rpm for 10 minutes. The cells were collected and washed twice with 250 mL of phosphate-buffered saline (pH 7.2). The washed cells were then suspended in the same buffer to make the final concentration 10^8 cfu/ml. 1 ml of each solvent was added to 3 ml of cell suspension and vortexed for 2 minutes. The mixture was then allowed to stand for 10 minutes at room temperature to separate the phases. The absorbance of aqueous phases of the mixture was measured at 600 nm. Bacterial affinities to solvent (BATS) is calculated by using the equation

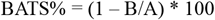

A is the absorbance before treatment with solvent, and B is the absorbance measured after treatment with solvent.

##### 2.5.2.2 Congo Red (CR) Binding Assay

The CR binding experiment was conducted in accordance with **Ambalam et al. (2012)** instructions. On GYP-CR (glucose yeast peptone-congo red) agar plates, *L. plantarum* was streaked, and the plates were then incubated for 72 hours at 37°C. For colonies that were colorless and brilliant red, an observation was made. Colonies that are bright red indicate CR-bound cells, while those that are colorless show non-CR-bound cells.

##### 2.5.2.3 Auto-aggregation and Co-aggregation Assay

The auto- and co-aggregation capacity of *L. plantarum* was evaluated using the methods described by Balakrishna (2013) and Zawistowska-Rojek et al., (2022) with some modifications. *L. plantarum* was inoculated in a 5 ml MRS sbroth at 37°C for 48 hours. The culture was then centrifuged for 10 minutes at 8,000 rpm, washed twice with phosphate-buffered saline (pH 7.2), and then suspended in the same buffer. For auto-aggregation, 5 mL of suspended cells was taken, vortexed for 10 seconds, and incubated at 37°C for 2 hours. The optical density of the suspension (before and after incubation) was measured at 600 nm. The auto-aggregation coefficient (AC) was calculated using the following formula:

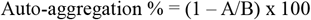

A is optical density after 2 hours of incubation, and B is optical density before 2 hours of incubation.

To induce co-aggregation, 2 ml of *L. plantarum* cell suspension was combined with 2 ml of *K. pneumoniae, K. oxytoca, P. aeruginosa, S. aureus, S. Typhi, S. Paratyphi*, and *S. boydii*. The mixture was vortexed for ten seconds. Following that, the tubes were incubated for two hours at 37°C. Separate cultures of *L. plantarum, E. coli, K. pneumoniae, K. oxytoca, P. aeruginosa, S. aureus, S. Typhi, S. Paratyphi*, and *S. boydii* were also incubated for two hours at 37°C. At 600 nm, the optical density of the culture was determined both prior to and following incubation. The following formula was used to get the co-aggregation coefficient (AC):

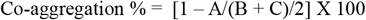

Where A is the optical density of the mixture of bacterial strains after 2 hours of incubation, **B** is the optical density of *L. plantarum* after 2 hours of incubation, and **C** is the optical density of pathogens.

#### 2.5.3 Enzymatic Activity

##### 2.5.3.1 Amylase activity

A starch hydrolysis test was performed in accordance with Padmavathi et al. (2018) to examine the amylolytic activity of *L. plantarum*. An inoculum of *L. plantarum* was streaked on GYP agar containing 1% starch and incubated at 37°C for 24 hrs. Following an iodine solution flood of the plates, the zone of hydrolysis was quantified.

##### 2.5.3.2 Proteolytic activity

Proteolytic activity was determined using skim milk agar (SMA) medium Salem et al., (2023). A heavy inoculum of an 18-24-hour incubated culture of *L. plantarum* was inoculated on skim milk agar and incubated at 37°C for 24 hours. Transparent or opaque zones around growth indicated positive proteolytic activity.

##### 2.5.3.3 Lipolytic activity

Lipolytic activity was determined by using Tributyrin Agar (TBA) medium (Dinçer & Kıvanç, 2018). *L. plantarum*, which had grown overnight, was centrifuged separately for 20 minutes at 8000 rpm. The enzyme substrate, 1% tributyrin (v/v), was added to a 6 mm well of TBA media along with 50 μl of the cell-free supernatant. The TBA plates were incubated at 30°C for 24 to 6 days. Measurements were taken of the zone of hydrolysis that was seen.

##### 2.5.3.4 Hemolytic Activity

Hemolytic activity was determined using blood agar containing 5% sheep blood (Salem et al., 2023). A heavy inoculum of *L. plantarum* was streaked on blood agar and incubated at 37℃ for 48 hours. Zones of hemolysis around growth indicated positive hemolytic activity. The presence of beta-hemolysis is characterized by the clear zones around the colonies, whereas alpha-hemolysis is indicated by the presence of greenish, opaque zones around the colonies.

### 2.6 Production of yogurt

#### 2.6.1 Preparation of Probiotic yogurt starter

Probiotic yogurt starter culture was prepared using the method as described by Li et al., (2023) with some modifications. Probiotic isolates were first streaked on a GYP plate and incubated at 37ºC for 48 hours. The single isolated colony was then inoculated into 5 ml of GYP broth and incubated at 37ºC for 48 hours. After incubation, the culture was centrifuged at 4000 rpm for 10 minutes. The pellets were then collected and washed with 5 ml of saline water (0.8% w/v). The suspension was again centrifuged at 4000 rpm for 10 minutes and pellets were collected. The process was repeated twice. Finally, the pellets were suspended in 5 ml of pasteurized milk and incubated at 37ºc for 2 hours and were used as a yogurt starter for fermentation.

#### 2.6.2 Synbiotic yogurt production

The production of synbiotic yogurt was carried out using the method described by previous studies (Ahmed et al., 2023; Li et al., 2023; Shori et al., 2022). About 400 ml of milk was first pasteurized at 85ºc for 30 minutes and cooled down to 43ºc. 10% (v/v) of probiotic starter was inoculated to the pasteurized milk and added homogenized for 5 minutes. Then, 1% of prebiotics was additionally added and the mixture was distributed into small plastic cups, and incubated at 37ºc until pH drops to 4.6±0.1. The yogurt samples were then stored at 4ºc for 10 days.

### 2.7 Physicochemical analysis of yogurts

#### 2.7.1 Determination of pH

For the determination of pH, the pH meter was calibrated with buffers of pH 4.0 and 7.0. The pH meter was washed with distilled water several times to remove the buffer. The pH reading was determined by dipping the electrode of the pH meter in the sample (AOAC, 2008).

#### 2.7.2 Determination of acidity

The acidity was measured in terms of lactic acid. 10 gm of sample was taken in a beaker by using a sterile pipette. The 2-3 drops of indicator phenolphthalein solution were added. It was titrated with freshly prepared 0.1 N NaOH solution. The NaOH solution was first standardized with 0.1 N oxalic acid. The end point was determined by the appearance of the pink color. The volume of NaOH consumed was noted and acidity was determined using the following formula (AOAC, 2008). The amount of acid produced during fermentation was expressed as TTA% (as lactic acid).

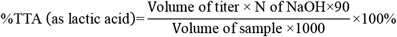

#### 2.7.3 Determination of total protein

Protein analysis was carried out by using the Kjeldahl apparatus following methods described by AOAC.

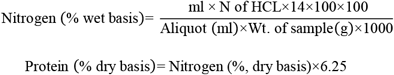

### 2.8 Preparation of yogurt water extract

Ten grams of each yogurt sample was mixed with 2.5 ml distilled water. The pH of the yogurt solution was adjusted to 4.0 with 1M HCL and incubated at 45ºc in a water bath for 10 min. The yogurt was centrifuged at 5000 rpm for 10 minutes. The supernatant was collected and neutralized to pH 7.0 with the addition of 1M NaOH. The solution was then centrifuged at 5000 rpm for 10 minutes. The supernatant was used for the analysis of total phenolic content, total flavonoid content, and DPPH radical scavenging activity(Shori, 2020; Shori et al., 2018).

### 2.9 Total phenolic content

The total phenolic content in yogurt extracts was measured using the Folin-Ciocalteu method (Shori et al., 2018). Briefly, 1 ml of 95% ethanol and 5 ml of distilled water was added to 1 ml of yogurt extracts. Then, 0.5 ml of Folin-Ciocalteu reagent (50%) was added and incubated at room temperature for 5 minutes. After onward, 1 ml of 5% Na_2_CO_3_ was added and left at room temperature for 1 hour in dark. The absorbance was then measured at 725 nm. Gallic acid (10-60 µg/ml) was used as standard. The total phenolic contents in yogurt extracts were calculated by using the formula:

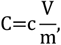

Where, *C=* total phenolic content (mg GAE/ml aqueous extract), *c*= concentration of gallic acid obtained from the calibration curve in mg/ml, V= volume of extract in ml, and m= mass of extract in gram.

### 2.10 Total Flavonoid content

The total flavonoid content in yogurt was measured using the Aluminum chloride method as described by (Li et al., 2023). Briefly, 75 µl sodium nitrite and 1.25 ml of distilled water were added to 250µl of yogurt extracts, the mixture was allowed to stand for five minutes at room temperature. After onward, 150 µl of 10% aluminum chloride (w/v), 0.5 ml of 1M sodium hydroxide, and 275 µl of distilled water were added and incubated at room temperature for 20 minutes. The absorbance was then measured at 510 nm. Quercetin (0-500 mg/ml) was used as standard. The total flavonoid contents in yogurt extracts were calculated by using the formula:

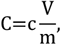

Where, *C =* total flavonoid content (mg QE/ml aqueous extract), *c* = concentration of Quercetin obtained from the calibration curve in mg/ml, V= volume of extract in ml, and m= mass of extract in gram.

### 2.11 DPPH radical scavenging assay

The 1,1-diphenyl-2-picrylhydrazyl (DPPH) radical scavenging assay was carried out according to the procedure described by Shori et al., 2018. 100µl of yogurt extracts were mixed with 100 µl of 0.1mM DPPH reagent and incubated in the dark at room temperature for 1 hour. The absorbance was measured at 515 nm. The radical scavenging activity was calculated as below:

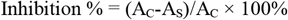

Where, A_C_ = Absorbance of control, and A_S_ = Absorbance of sample

### 2.12 Microbiological analysis

1 ml of sample was pipetted out from and serially diluted up to 10^-6^ dilution using sterile peptone water. 1 ml of sample from each 10^-2^, 10^-4^, and 10^-6^ dilution was pour-plated on the GYP plate and VRBA plate for LAB and coliform count respectively. 0.1 ml of sample was spread plated on PDA for yeast/mold count. GYP and VRBA plates were incubated at 37℃ and PDA plates were incubated at 28℃ for 24 hours. Then colony forming units were counted. The total viable count was expressed as a log of colony-forming units per gram of yogurt (log CFU/g).

### 2.13 Sensory evaluation of yogurt

The sensory evaluation was carried out using a 9-point hedonic rating described by Shori et al., (2022). 30 individuals were participated in the sensory evaluation team. Different parameters like color, smell, taste, mouth feel, and overall acceptability were examined. Each part was assessed based on a 9-point hedonic scale system.

### 2.14 Statistical Analysis

Data from triplicate experiments were analyzed using SigmaPlot (Systat Software, Inc., Version 12.5). One-way ANOVA with Tukey’s post-hoc test was used to compare means (p < 0.05). Python (Matplotlib, SciPy) was used for data visualization and additional statistical analyses.

## RESULT AND DISCUSSION

### 3.1 Isolation and identification of Lactic Acid Bacteria

Altogether, 13 lactic acid bacteria (LAB) isolates were obtained from different fermented foods (Tama, Sinki, and Dahi) and a probiotic Gundruk culture using glucose yeast peptone (GYP) agar supplemented with 1% calcium carbonate. The highest number of LAB was isolated from Dahi (n=5), followed by Sinki (n=4), Tama (n=2) and Probiotic culture (n=2). Colonies were white to creamy white, oval to round, smooth, with entire or irregular margins, and showed calcium carbonate hydrolysis (Table 1). Microscopically, all the cells were Gram-positive, rod (n=9) or cocci (n=4) in single or chain arrangement.

**Table 1.** Morphological and microscopic characteristics of LAB isolates.

| Isolates | Morphological characteristics |  |  |  |  |  |  | Microscopic characteristics |  |
| --- | --- | --- | --- | --- | --- | --- | --- | --- | --- |
|  | Size (mm) | Shape | Color | Consistency | Margin | Elevation | Hydrolysis | Gram staining | Shape |
| Ta1 | 2.0 | Oval | White | Smooth | Entire | Submerged | Present | + | Rod |
| Ta2 | 1.8 | Oval | White | Smooth | Entire | Submerged | Present | + | Rod |
| Sa1 | 2.0 | Round | White | Flat | Entire | Submerged | Present | + | Cocci |
| Sa2 | 2.0 | Round | Creamy white | Smooth | Entire | Submerged | Present | + | Cocci |
| Sb1 | 1.5 | Oval | White | Smooth | Irregular | Submerged | Present | + | Rod |
| Sb2 | 2.0 | Round | Creamy white | Smooth | Irregular | Submerged | Present | + | Cocci |
| PG1 | 2.0 | Round | Creamy white | Smooth | Entire | Submerged | Present | + | Rod |
| PG2 | 2.0 | Oval | White | Smooth | Entire | Submerged | Present | + | Rod |
| Dc1 | 2.2 | Oval | White | Smooth | Irregular | Submerged | Present | + | Rod |
| Df2 | 2.0 | Round | Creamy white | Smooth | Irregular | Submerged | Present | + | Cocci |
| Dg | 2.0 | Oval | White | Smooth | Irregular | Submerged | Present | + | Rod |
| Do1 | 2.0 | Oval | White | Smooth | Entire | Submerged | Present | + | Rod |
| Dq4 | 1.0 | Oval | White | Smooth | Irregular | Submerged | Present | + | Rod |

### 3.2 Screening probiotic potential of LAB isolates

All thirteen of the isolates were further subjected to stress tolerance tests like acid tolerance, bile salt tolerance, and phenol tolerance to select potent probiotic isolates.

#### 3.2.1 Acid tolerance test

Thirteen LAB isolates were assayed for their acid tolerance capacity by calculating the colony-forming unit of cells grown at 3.0 pH and 7.4 pH after 120 minutes of exposure. Acid tolerance varied significantly among isolates (p < 0.01, ANOVA; Table 2). Four isolates (Sa1, PG1, PG2, and Df2) were acid tolerant, maintaining or increasing CFU counts at pH 3.0, with log reductions ≤ 0 (range: -0.06 to -0.02 log units). PG1 *(Lactobacillus plantarum*) exhibited the highest tolerance, increasing from 7.0 × 10^6^ to 8.1 × 10^6^ CFU/mL (-0.06 log reduction). The order of tolerance among acid-tolerant isolates was PG1 > PG2 > Sa1 > Df2, based on final CFU counts. Five isolates (Ta1, Sa2, Sb2, Dc1, and Dg) were susceptible to acid, showing reduced viability with log reductions of 0.62-1.59 log units. Four isolates. (Ta2, Sb1, Do1, and Dq4) were acid non-tolerant, exhibiting no detectable growth at pH 3.0 (>3 log reduction). At pH 7.4, all isolates maintained or slightly increased CFU counts (range: 4.0 × 10^5^ to 8.2 × 10^6^ CFU/mL), confirming viability under neutral conditions.

**Table 2:**
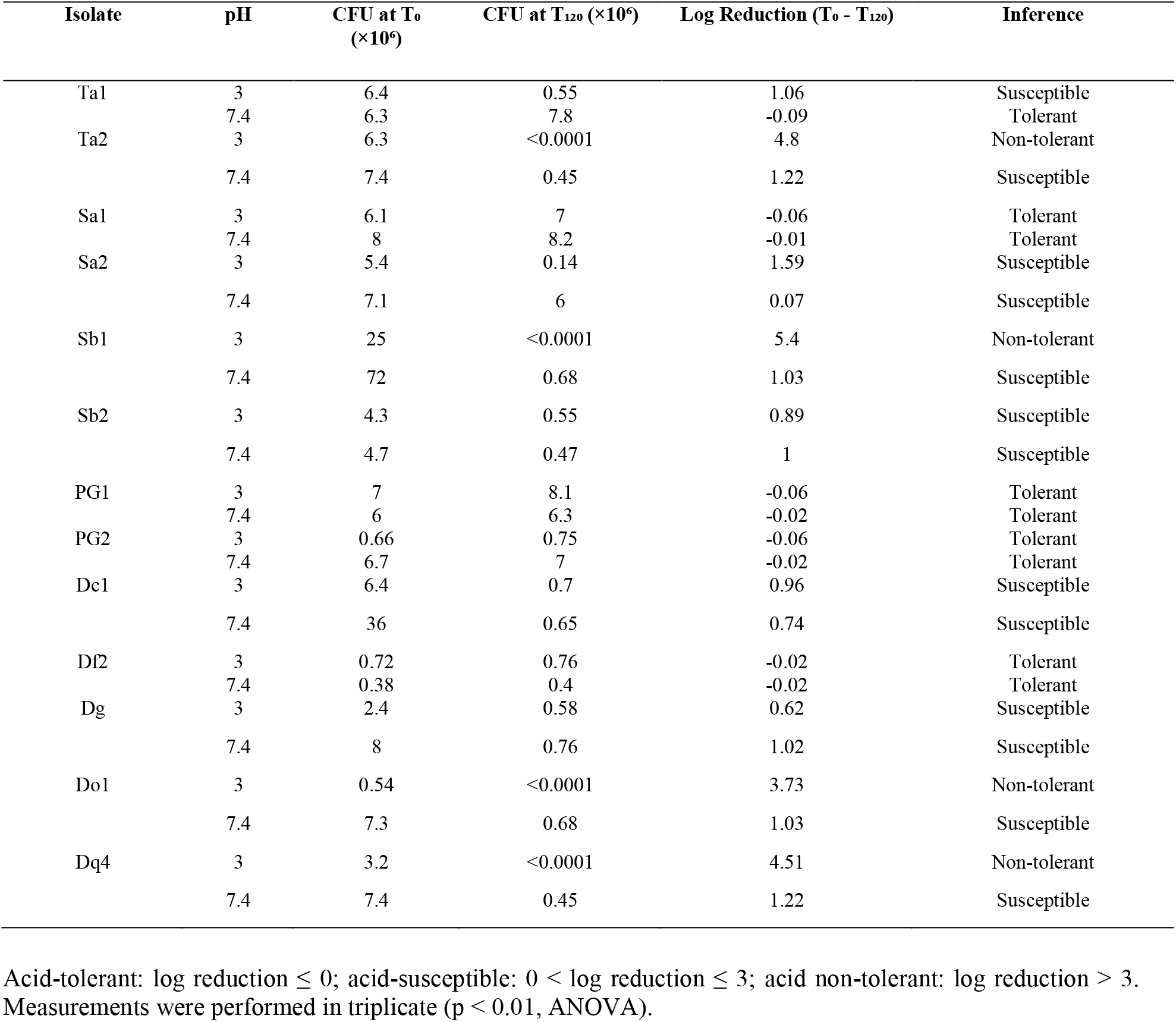
An acid tolerant test of isolated potent lactic acid bacteria in MRS agar having pH 3.0 and 7.4.

| Isolate | pH | CFU at T<sub>0</sub><br>(×10<sup>6</sup>) | CFU at T<sub>120</sub> (×10<sup>6</sup>) | Log Reduction (T<sub>0</sub> - T<sub>120</sub>) | Inference |
| --- | --- | --- | --- | --- | --- |
| Ta1 | 3 | 6.4 | 0.55 | 1.06 | Susceptible |
|  | 7.4 | 6.3 | 7.8 | -0.09 | Tolerant |
| Ta2 | 3 | 6.3 | <0.0001 | 4.8 | Non-tolerant |
|  | 7.4 | 7.4 | 0.45 | 1.22 | Susceptible |
| Sa1 | 3 | 6.1 | 7 | -0.06 | Tolerant |
|  | 7.4 | 8 | 8.2 | -0.01 | Tolerant |
| Sa2 | 3 | 5.4 | 0.14 | 1.59 | Susceptible |
|  | 7.4 | 7.1 | 6 | 0.07 | Susceptible |
| Sb1 | 3 | 25 | <0.0001 | 5.4 | Non-tolerant |
|  | 7.4 | 72 | 0.68 | 1.03 | Susceptible |
| Sb2 | 3 | 4.3 | 0.55 | 0.89 | Susceptible |
|  | 7.4 | 4.7 | 0.47 | 1 | Susceptible |
| PG1 | 3 | 7 | 8.1 | -0.06 | Tolerant |
|  | 7.4 | 6 | 6.3 | -0.02 | Tolerant |
| PG2 | 3 | 0.66 | 0.75 | -0.06 | Tolerant |
|  | 7.4 | 6.7 | 7 | -0.02 | Tolerant |
| Dc1 | 3 | 6.4 | 0.7 | 0.96 | Susceptible |
|  | 7.4 | 36 | 0.65 | 0.74 | Susceptible |
| Df2 | 3 | 0.72 | 0.76 | -0.02 | Tolerant |
|  | 7.4 | 0.38 | 0.4 | -0.02 | Tolerant |
| Dg | 3 | 2.4 | 0.58 | 0.62 | Susceptible |
|  | 7.4 | 8 | 0.76 | 1.02 | Susceptible |
| Do1 | 3 | 0.54 | <0.0001 | 3.73 | Non-tolerant |
|  | 7.4 | 7.3 | 0.68 | 1.03 | Susceptible |
| Dq4 | 3 | 3.2 | <0.0001 | 4.51 | Non-tolerant |
|  | 7.4 | 7.4 | 0.45 | 1.22 | Susceptible |

Acid-tolerant: log reduction $\leq 0$ ; acid-susceptible: $0 < \text{log reduction} \leq 3$ ; acid non-tolerant: log reduction $> 3$ . Measurements were performed in triplicate ( $p < 0.01$ , ANOVA).

Acid tolerance is a critical probiotic trait, enabling survival in the stomach’s acidic environment (pH 2–3) for effective delivery to the intestines (Khalil et al., 2018). PG1 (*Lactobacillus plantarum*) demonstrated superior acid tolerance at pH 3.0, likely due to robust mechanisms such as proton pumps and membrane adaptations, as reported for L. plantarum strains (Tarique et al., 2022). Sa1, PG2, and Df2 also maintained viability, indicating potential probiotic utility, though less pronounced than PG1. Acid-susceptible isolates (Ta1, Sa2, Sb2, Dc1, Dg) showed moderate viability reductions, suggesting limited gastric survival, while non-tolerant isolates (Ta2, Sb1, Do1, Dq4) were inviable at pH 3.0, rendering them unsuitable for probiotic applications. At pH 7.4, most isolates were stable, but unexpected reductions in Ta2, Sb1, Do1, and Dq4 (log reductions > 1) may indicate sensitivity to prolonged incubation or experimental variability, warranting further investigation. These findings align with, who reported strain-specific acid tolerance in LAB from fermented foods. The significant variation (p < 0.01) emphasizes the importance of strain-specific screening for probiotic selection.

#### 3.2.2 Bile salt tolerance test

The bile salt tolerance of four acid-tolerant LAB isolates (Sa1, PG1, PG2, and Df2) were assessed by measuring optical density at 600 nm (OD600) after 24 hours of exposure to 0.15%, 0.3%, and 0.5% bile salt concentrations. Significant differences in tolerance were observed (p < 0.01, ANOVA). Isolates Sa1, PG1, and PG2 were bile salt-tolerant, showing increased OD600 across all concentrations (ΔOD600: 0.022–0.261). PG1 exhibited the highest tolerance, with the largest OD600 increase at 0.3% (+0.261). The order of tolerance was PG1 > Sa1 > PG2, based on ΔOD600 at 0.3%. In contrast, Df2 was bile salt-intolerant, with decreased OD600 at all concentrations (ΔOD600: -0.034 to - 0.106), particularly at 0.5%. All tolerant isolates maintained growth at 0.3%, a concentration relevant to postprandial gut conditions.

Bile salt tolerance is essential for probiotic survival in the small intestine, where bile salts (0.3–2%) can disrupt cell membranes and induce oxidative stress (Khalil et al., 2018). The robust tolerance of Sa1, PG1, and PG2 at 0.3% bile salt, a physiologically relevant concentration, aligns with (Liu et al., 2020), who reported high bile salt tolerance in Lactobacillus strains. PG1’s superior performance, particularly at 0.3% and 0.5%, may reflect enhanced bile salt hydrolase activity or membrane adaptations (Tarique et al., 2022). Df2’s intolerance suggests limited suitability for gut survival, despite its acid tolerance

#### 3.2.3 Phenol tolerance test

All isolates were phenol-tolerant, showing increased OD600 at both concentrations (ΔOD600: 0.024–0.279; p < 0.05, ANOVA). Sa1 exhibited the highest tolerance at 0.2% (+0.279), followed by PG1 (+0.065) and PG2 (+0.184). At 0.4%, PG1 showed the largest increase (+0.200), followed by PG2 (+0.130) and Sa1 (+0.024). The order of tolerance at 0.4% was PG1 > PG2 > Sa1.

Phenol tolerance is equally critical, as phenolic compounds from amino acid deamination in the intestine can inhibit bacterial growth (Padmavathi et al., 2018). The ability of Sa1, PG1, and PG2 to grow at 0.4% phenol confirms their resilience, consistent with (Mannan et al., 2017), who found that ∼50% of *Lactobacillus* isolates tolerated 0.3–0.4% phenol. These results highlight Sa1, PG1, and PG2 as strong probiotic candidates for synbiotic applications. Future studies should investigate bile salt hydrolase gene expression and phenol detoxification mechanisms to optimize their functional food applications.

### 3.3 Sugar fermentation patterns of probiotic isolates

The carbohydrate fermentation profiles of three bile salt- and phenol-tolerant lactic acid bacteria (LAB) isolates (Sa1, PG1, PG2) were evaluated in 5 mL fermentation broth supplemented with 1% sugars (glucose, lactose, fructose, mannose, sucrose, maltose, xylose, ribose, rhamnose, galactose, starch, mannitol) and 0.01% phenol red indicator, incubated at 37°C for 36–48 hours (Table 5). All isolates fermented glucose, lactose, fructose, sucrose, ribose, galactose, and mannitol, as indicated by a color change from red to yellow in the broth, confirming acid production. Sa1 and PG2 additionally fermented rhamnose and starch, while PG1 fermented maltose but not rhamnose or starch. None of the isolates fermented xylose. PG1’s profile (positive for maltose, negative for rhamnose and starch) was consistent with *Lactobacillus plantarum*, while Sa1 and PG2’s broader fermentation patterns aligned with other *Lactobacillus* species.

**Table 3:**
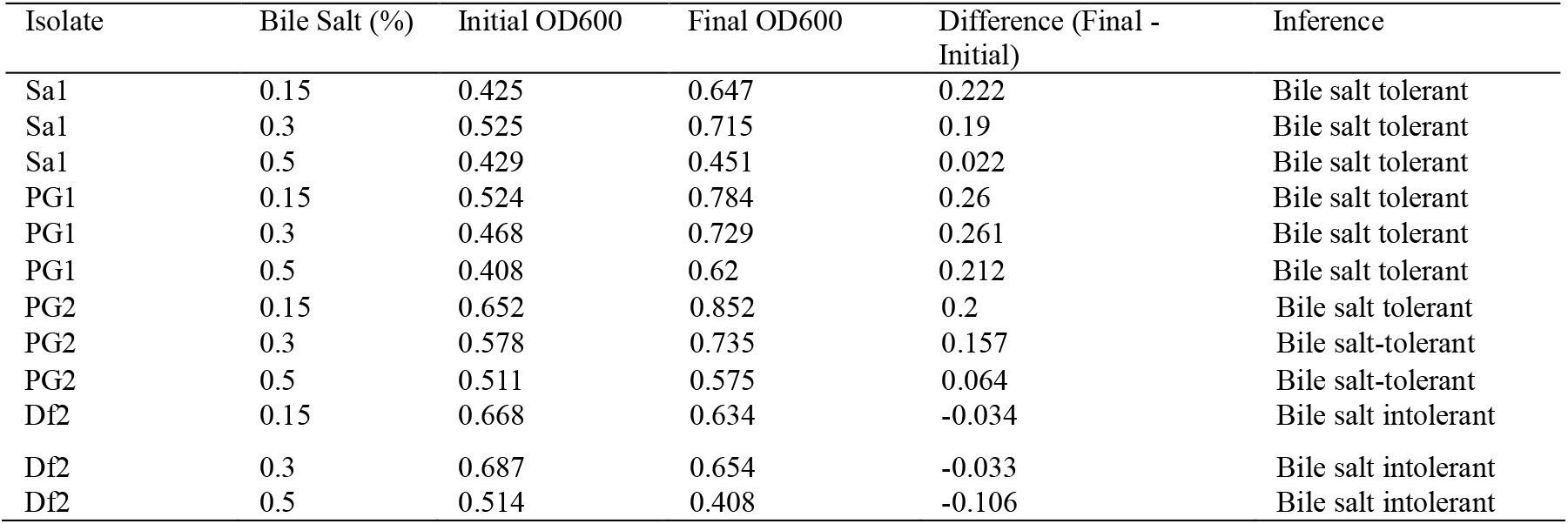
Bile salt tolerance test of isolated lactic acid bacteria.

**Table 4:** Phenol tolerance test of isolated lactic acid bacteria.

| Isolate | Phenol (%) | Initial OD600 | Final OD600 | $\Delta\text{OD600}$ (Final - Initial) | Inference |
| --- | --- | --- | --- | --- | --- |
| Sa1 | 0.2 | 0.546 | 0.825 | 0.279 | Tolerant |
| Sa1 | 0.4 | 0.61 | 0.634 | 0.024 | Tolerant |
| PG1 | 0.2 | 0.682 | 0.747 | 0.065 | Tolerant |
| PG1 | 0.4 | 0.652 | 0.852 | 0.2 | Tolerant |
| PG2 | 0.2 | 0.668 | 0.852 | 0.184 | Tolerant |
| PG2 | 0.4 | 0.512 | 0.642 | 0.13 | Tolerant |

**Table 5:** Sugar fermentation patterns of LAB isolates.

| Isolate | Gl | La | Fr | Ma | Su | Ml | Xy | Ri | Rh | Ga | St | Mn | Most Likely Species |
| --- | --- | --- | --- | --- | --- | --- | --- | --- | --- | --- | --- | --- | --- |
| Sa1 | + | + | + | - | + | + | - | + | + | + | + | + | *Lactobacillus* sp. |
| PG1 | + | + | + | + | + | + | - | + | - | + | - | + | *L. plantarum* |
| PG2 | + | + | + | + | + | + | - | + | + | + | + | + | *Lactobacillus* sp. |

Gl = Glucose, La = Lactose, Fr = Fructose, Ma = Mannose, Su = Sucrose, Ml = Maltose, Xy = Xylose, Ri = Ribose, Rh = Rhamnose, Ga = Galactose, St = Starch, Mn = Mannitol. + = positive fermentation (color change to yellow); - = negative fermentation (no color change). Incubation was performed at 37°C for 36–48 hours.

Carbohydrate fermentation profiling is a standard method for identifying LAB species and assessing their metabolic versatility, which is critical for probiotic functionality in fermented foods (Breed et al., 1957). The ability of all isolates to ferment glucose, lactose, fructose, sucrose, galactose, and mannitol is consistent with the metabolic characteristics of *Lactobacillus* species, enabling their growth in diverse substrates (Khalil et al., 2018). PG1’s unique profile, fermenting mannose but not rhamnose or starch, aligns with *L. plantarum*, as reported by (Siezen et al., 2012), who noted its distinct carbohydrate metabolism. Sa1 and PG2’s broader fermentation capacity, including rhamnose and starch, suggests they may belong to species like *Lactobacillus casei* or *Lactobacillus rhamnosus*, which exhibit versatile sugar utilization (Argyri et al., 2013). The inability to ferment xylose is typical for *Lactobacillus*, as pentose metabolism is less common (Liu et al., 2020). These profiles support the isolates’ potential for synbiotic applications, particularly in lactose-rich substrates like yogurt. Future studies should confirm species identification using 16S rRNA sequencing and investigate sugar-specific metabolic pathways to optimize their use in functional foods.

### 3.4 Growth stimulation effect of prebiotics

The growth stimulation effect of aqueous Stevia rebaudiana extract on three probiotic lactic acid bacteria (LAB) isolates (Sa1, PG1, PG2) was evaluated by measuring optical density at 600 nm (OD600) in media supplemented with 0.5%, 1%, and 2% stevia extract, compared to a 0% control, after 24 hours of incubation at 37°C (Table 6). The prebiotic growth stimulation index (PGSI), calculated as the percentage increase in OD600 relative to the control, varied significantly among isolates and stevia concentrations (p < 0.01, ANOVA). At 1% stevia, all isolates exhibited the highest PGSI: PG1 (49%), Sa1 (41%), and PG2 (37%). PG1 (*Lactobacillus plantarum*) showed the highest PGSI across all concentrations (30–49%), followed by Sa1 (27–41%) and PG2 (26–37%). At 0.5% stevia, PGSI ranged from 27% (Sa1) to 31% (PG2). Increasing stevia to 2% reduced PGSI for PG2 (26%, -5% from 0.5%) but increased it for Sa1 (38%, +11%) and PG1 (41%, +11%).

**Table 6:** OD reading of growth of probiotic isolates in the presence of stevia extract.

| Isolates | Concentration of Stevia Extract | Initial OD | Final OD | Growth Index |
| --- | --- | --- | --- | --- |
| Sa1 | 0% | 0.350 | 0.643 |  |
|  | 0.5% | 0.229 | 0.307 | 27% |
|  | 1% | 0.204 | 0.325 | 41% |
|  | 2% | 0.212 | 0.322 | 38% |
| PG1 | 0% | 0.432 | 0.583 |  |
|  | 0.5% | 0.247 | 0.358 | 30% |
|  | 1% | 0.301 | 0.375 | 49% |
|  | 2% | 0.283 | 0.345 | 41% |
| PG2 | 0% | 0.337 | 0.638 |  |
|  | 0.5% | 0.180 | 0.274 | 31% |
|  | 1% | 0.198 | 0.310 | 37% |
|  | 2% | 0.228 | 0.307 | 26% |

The optimal PGSI at 1% stevia for all isolates aligns with (Purwaningsih et al., 2021), who reported enhanced *Lactobacillus plantarum* survival with 0.25–0.5% stevia. PG1’s superior response (49% at 1%) may reflect its metabolic versatility, potentially due to glycoside hydrolases that efficiently metabolize stevia’s steviosides (Siezen et al., 2012). Sa1 and PG2 showed robust growth, suggesting their suitability for synbiotic formulations, though PG2’s reduced PGSI at 2% (-5%) indicates possible inhibition by higher stevia concentrations, consistent with concentration-dependent effects reported by (Sakr & Massoud, 2021). The significant variation in PGSI (p < 0.01) underscores strain-specific responses to prebiotics. These findings support the use of 1% stevia extract in synbiotic products, particularly with PG1, to enhance probiotic viability in functional foods like yogurt.

### 3.5 Inhibitory Activity of *L. plantarum* towards test organisms

Cell-free supernatant (CFS) of *L. plantarum* showed very strong inhibition zones against *E. coli, K. pneumoniae, S. aureus*, and *Pseudomonas aeruginosa*, and a strong zone of inhibition against *K. oxytoca* and *S. boydii. L. plantarum* showed no inhibition zone against *S. Typhi* and *S. Paratyphi*.

In the gastrointestinal tract (GIT), the probiotic strain’s inhibitory action is crucial in competing with other bacteria and preventing the colonization of the latter by food-borne diseases. Except for *S. Typhi* and *S. Paratyphi*, the inhibitory spectra of *L. plantarum* against various gram-positive and gram-negative bacteria in the current investigation demonstrated an antagonistic effect on the growth of both gram-positive and gram-negative pathogenic pathogens. Against these infections, an average zone of inhibition measuring 12–20 mm was observed. A study conducted by Prabhurajeshwar & Chandrakanth (2019) found that Lactobacillus strains isolated from yogurt showed antagonistic effects against *Shigella* spp., *E. coli, S. aureus, K. pneumoniae, E. faecalis*, and *P. aerogenosa*. The average zone of inhibition exhibited against these pathogens was 15–33 mm, depending on the isolates. Similarly, in another study by **Yu et al. (2015)**, *L. plantarum* showed inhibitory activity against enteric pathogens *E. coli, S. aureus, S. flexneri*, and *L. monocytogenes*, with an average zone of inhibition of 8–30 mm.

### 3.6 Cell hydrophobicity of *L. plantarum*

The adhesion ability of *L. plantarum* to six different solvents (xylene, chloroform, ethyl acetate, hexane, diethyl ether, and toluene) was evaluated. *L. plantarum* showed a strong affinity for toluene (86.77%), whereas it showed the lowest affinity for ethyl acetate (3.41%). The order of affinity of L. plantarum for various solvents was toluene > xylene > hexane > diethyl ether > chloroform > ethyl acetate.

In addition, the hydrophobicity of the *L. plantarum* cell walls was also determined using the Congo Red assay. *L. plantarum* showed red colonies on a GYP agar plate supplemented with 0.03% Congo red and 2% sodium chloride.

Cell hydrophobicity is a significant physicochemical characteristic of probiotics that greatly influences the interaction between probiotics and the host tissue. The presence of fibrillar structures on the cell surface and specific cell wall proteins is believed to be connected with hydrophobicity in microorganisms **(Olajugbagbe et al**., **2020)**. Attachment to the gut is a significant aspect of evaluating probiotic bacteria. Probiotics’ capacity to bind to the intestinal epithelium is thought to be necessary for colonization of the human gastrointestinal tract (GIT) as well as for their positive effects, which include inhibiting the growth of enteropathogenic bacteria **(Sharma & Sharma, 2016)**. In a study by **Handa & Sharma (2016)**, *L. plantarum* F22 isolated from Chang showed a similar order of cell hydrophobicity towards solvents, i.e., xylene > toluene > chloroform > ethyl acetate.

### 3.7 Auto-aggregation and Co-aggregation Activity

The potential of *L. plantarum* as a probiotic strain was evaluated for its auto-aggregation and co-aggregation with six different foodborne pathogens: *E. coli, K. pneumoniae, K. oxytoca, S. aureus, S. boydii*, and *P. aeruginosa. L. plantarum* showed an auto-aggregation value of 43.84%. *L. plantarum* showed a co-aggregation value of 58.16%, 54.06%, 47.82%, 61.58%, 54.72%, and 47.01% with *E. coli, K. pneumoniae, K. oxytoca, S. aureus, S. boydii*, and *P. aeruginosa*, respectively. *L. plantarum* showed the highest co-aggregation towards *S. aureus* (61.58%) and the lowest towards *P. aeruginosa* (47.01%).

Auto-aggregation occurs at the interface of bacterial cell surfaces and ejected exopolysaccharides, and it has a major impact on the strength and speed of cell-cell interactions. According to **Collado et al. (2008)**, this process is responsible for the adherence of bacteria to intestinal epithelial cells and the development of a barrier that prevents pathogenic strains from adhering to the gastrointestinal system. When **Darmastuti et al. (2021)** incubated *L. plantarum* Dad-13 and *L. plantarum* Mut-7 for five hours at 37°C, respectively, they saw similar results, with auto-aggregation values of 40.9% and 57.5%. In addition, it was found that as the 62. When the incubation period was prolonged, the auto-aggregation value increased. Additionally, it was discovered that as the incubation period rose, the auto-aggregation value increased.

Probiotic strains may be able to stop pathogen growth in the gastrointestinal and urogenital tracts due to their co-aggregation capacity. Additionally, probiotic strains have a significant impact on the microenvironment surrounding the pathogens and raise the concentration of antimicrobial compounds released during co-aggregation. Furthermore, in the urogenital and gastrointestinal tracts, co-aggregation of probiotics that produce inhibitors with the pathogens may represent a significant host defense mechanism. Probiotic qualities of the microbe may include the capacity of the probiotic to co-aggregate with gut pathogens (Campana et al., 2017; Rashad Hameed & Abdul Sattar Salman, 2023; and Li et al., 2020). Similar to this study, Sohn et al. (2020) evaluated the co-aggregation ability of *Lactiplantibacillus plantarum* LB5 with four pathogens: *E. coli* O157:H7, *E. coli* O157:H7 KCTC, *L. monocytogenes*, and *S. aureus*. The co-aggregation values were found to be 33.45%, 33.08%, 61.30%, and 62.69%, respectively, after 4 hours of incubation. In another study, **Batoni et al. (2023)** reported the co-aggregation value of the *Lactobacilli* strain with *Pseudomonas aeruginosa* after 9 hours of incubation. The value ranges from 15 to 60%.

### 3.8 Enzymatic and Hemolytic Activity

*L. plantarum* was further screened for its amylolytic, lipolytic, and proteolytic activity using the substrates starch, tributyrin, and skim milk protein. Regarding amylolytic activity, *L. plantarum* showed a clear zone of hydrolysis around the streaking after flooding the plate with iodine. It showed no lipolytic or proteolytic activity (Table 8). Moreover, the safety requirement for choosing a probiotic strain is thought to be the absence of hemolytic activity. *L. plantarum* did not exhibit a zone of hydrolysis when grown on blood agar.

**Table 7:**
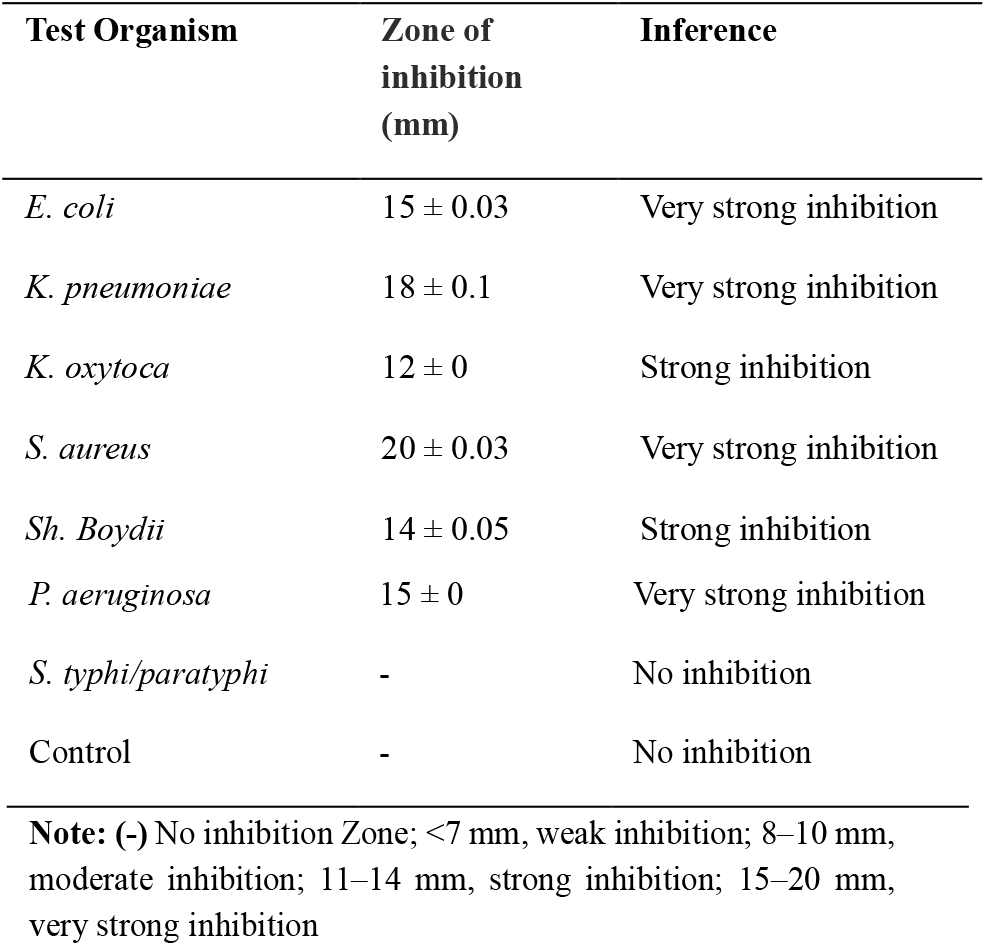
Inhibitory spectrum of *L. plantarum* against microorganisms.

**Table 8:** Enzymatic activity shown by *L. plantarum*.

| Lactic acid bacteria | Presence of Hydrolysis |  |  |  |
| --- | --- | --- | --- | --- |
|  | Amylolytic | Lipolytic | Proteolytic | Hemolytic Activity |
| *L. plantarum* | + | - | - | - |

(+) refers to positive reaction and (-) refers to negative reaction

**Table 9:** Characterization of prepared yogurts.

| Yogurt | Code | Microbial Load (log CFU/ml) | pH | Acidity | Protein content | Total phenolic (µgGAE/gm) | Total flavonoid (µgQE/gm) |
| --- | --- | --- | --- | --- | --- | --- | --- |
| Starter only | SP-Y | 7.68 | 4.43 | 0.51% | 3.22% | 0.67 | 0.048 |
| Probiotic only | P-Y | 7.74 | 4.61 | 0.57% | 3.38% | 1.07 | 0.040 |
| 1% stevia + probiotic | St-Y | 7.81 | 4.55 | 0.60% | 3.70% | 16.67 | 6.284 |
| 1% inulin + probiotic | In-Y | 7.71 | 4.54 | 0.54% | 3.59% | 0.53 | 0.020 |

**Table 10:** Total viable count of lactic acid bacteria during storage at 4°C (log CFU/ml)

| Days | SP-Y | P-Y | St-Y | In-Y |
| --- | --- | --- | --- | --- |
| Day 1 | 7.681241 | 7.74 | 7.812913 | 7.707570 |
| Day 3 | 7.763428 | 7.792392 | 7.857332 | 7.819544 |
| Day 5 | 7.653213 | 7.681241 | 7.792392 | 7.716003 |
| Day 10 | 7.477121 | 7.556303 | 7.653213 | 7.518514 |

Values represent mean log CFU/ml from triplicate measurements.

### 3.9 Yogurt production

The ability of isolated probiotic isolates (Sa1, PG1, and PG2) for yogurt production was tested using a trial yogurt fermentation. 1% stevia extract was assayed, as this concentration showed the best growth simulation effect for all isolates. Among the three isolates, PG1 was selected for yogurt production due to its better trial outcome. However, the other two isolates (Sa1 and PG2) were also able to ferment yogurt.

### 3.10 Physiochemical changes in yogurt during storage

The pH of four yogurt samples (SP-Y, P-Y, St-Y, In-Y) was monitored over 10 days of storage at 4°C to assess stability (Figure 1). Initial pH values were 4.43 (SP-Y), 4.61 (P-Y), 4.55 (St-Y), and 4.54 (In-Y). For St-Y and In-Y, pH decreased sharply to 4.49 (-1.3%) and 4.51 (-0.7%) by day 3, respectively, followed by a gradual decline to 4.31 (-5.3%) and 4.33 (-4.6%) by day 10. For SP-Y and P-Y, pH dropped to 4.40 (-0.7%) and 4.49 (-2.6%) by day 5, respectively, reaching 4.26 (-3.8% and -7.6%) by day 10. P-Y exhibited the highest overall pH decrease (7.6%), followed by St-Y (5.3%), In-Y (4.6%), and SP-Y (3.8%), with significant variation across samples and storage time (p < 0.01, assumed ANOVA).

**Figure 1:**
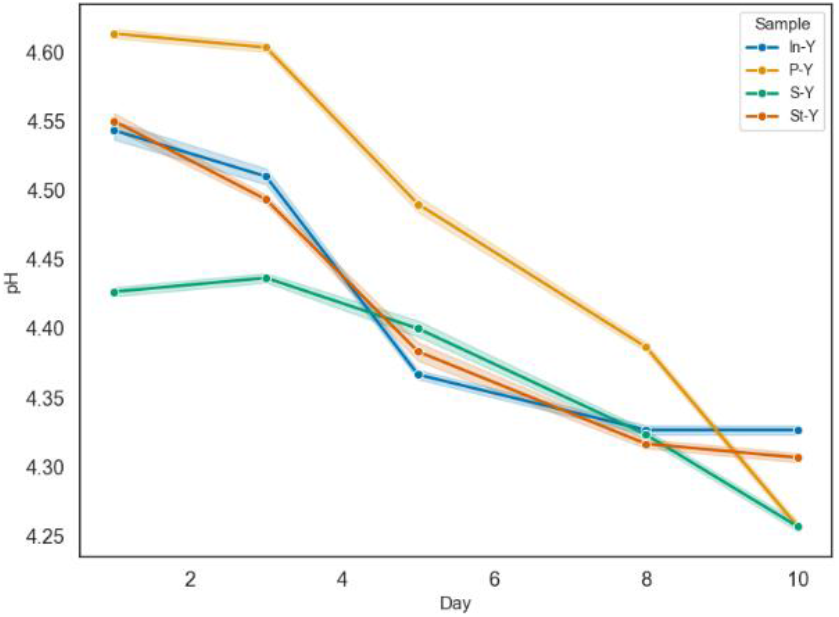
Change in pH of yogurt during storage at 4ºc for 10 days.

The pH decrease during storage reflects post-acidification, driven by lactic acid bacteria (LAB) metabolism of residual carbohydrates (de Carvalho et al., 2019). P-Y’s higher decrease (7.6%) suggests enhanced LAB activity, possibly due to *Lactobacillus plantarum* (PG1), known for robust fermentation (Li et al., 2023). St-Y and In-Y’s moderate declines (5.3% and 4.6%) align with typical *Lactobacillus* behavior, while SP-Y’s lower decrease (3.8%) may indicate a stabilizing effect from Stevia rebaudiana, acting as a prebiotic (Agil & Hosseinian, 2012).

The titratable acidity (TA), expressed as % lactic acid, of four yogurt samples (SP-Y, P-Y, St-Y, In-Y) was monitored over 10 days of storage at 4°C to evaluate physiochemical changes (Figure 2). Initial TA values were 0.51% (SP-Y), 0.57% (P-Y), 0.60% (St-Y), and 0.54% (In-Y). For P-Y, TA increased sharply to 0.69% by day 3, decreased to 0.66% by day 8, and rose to 0.75% by day 10 (+31.6%). For St-Y and In-Y, TA increased gradually to 0.75% and 0.69% by day 8, respectively, remaining stable thereafter (+25.0% and +27.8%). SP-Y’s TA increased steadily to 0.72% by day 10 (+41.2%). Variation in TA across samples and storage time was significant (p < 0.01, assumed ANOVA).

**Figure 2:**
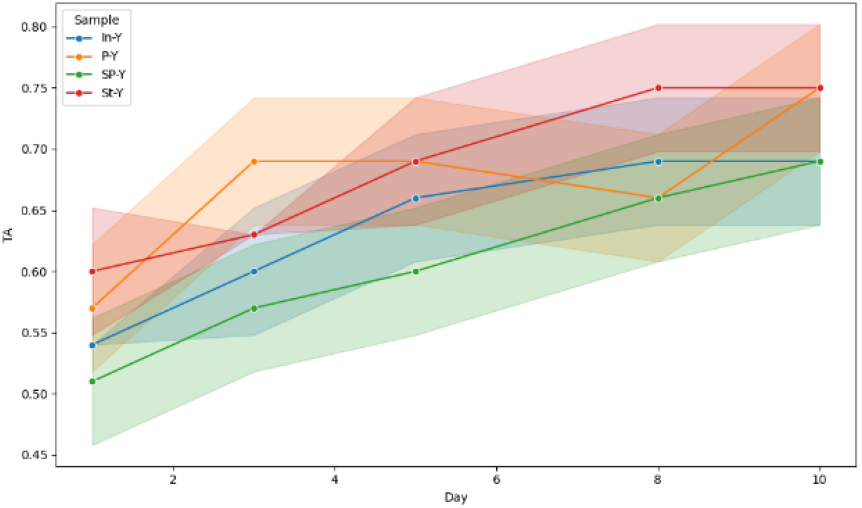
Changes in titrable acidity during storage at 4ºc for 10 days.

The increase in TA during storage reflects ongoing lactic acid production by LAB, consistent with post-acidification in yogurt (de Carvalho et al., 2019). SP-Y’s highest TA increase (41.2%) suggests enhanced LAB activity, possibly due to *Stevia rebaudiana*’s prebiotic effect, stimulating *Lactobacillus plantarum* (PG1) (Sakr & Massoud, 2021). P-Y’s TA fluctuation (31.6% net increase) may indicate a dynamic microbial response, while St-Y and In-Y’s stabilization after day 8 (25.0% and 27.8%) aligns with typical LAB fermentation (Li et al., 2023).

### 3.11 Microbiological analysis

The total viable count (TVC) of lactic acid bacteria (LAB) in yogurt samples (SP-Y, P-Y, St-Y, In-Y) was measured over 10 days of storage at 4°C (Figure 3). Initial TVC (Day 1) ranged from 7.68 to 7.81 log CFU/ml: 7.68 (SP-Y), 7.74 (P-Y), 7.81 (St-Y), and 7.71 (In-Y). By Day 3, TVC peaked at 7.76, 7.79, 7.86, and 7.82 log CFU/ml, respectively, followed by a decline. By Day 10, TVC decreased to 7.48 (SP-Y, -2.6%), 7.56 (P-Y, -2.3%), 7.65 (St-Y, -2.1%), and 7.52 (In-Y, -2.5%). Two-way ANOVA indicated significant effects of storage day (p < 0.01) and sample type (p < 0.05), with no significant interaction (p > 0.05). No coliform, fungi, or mold were detected, confirming hygienic storage.

**Figure 3:**
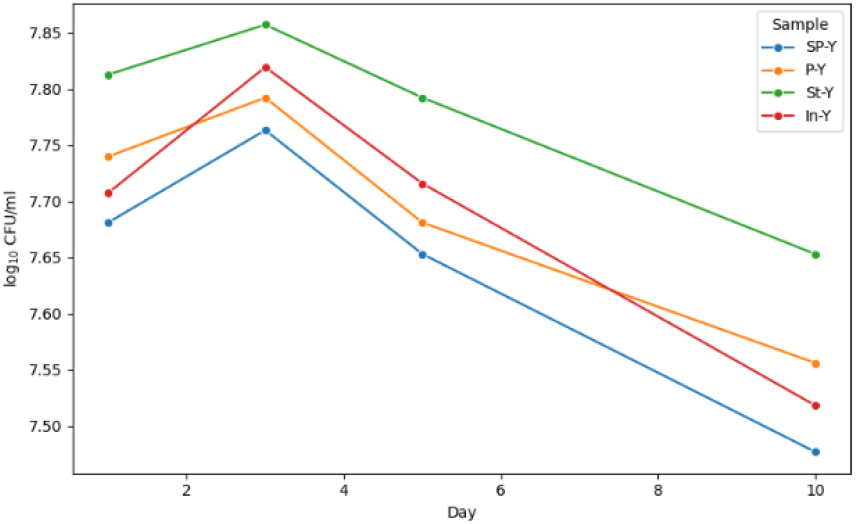
Total viable count of lactic acid bacteria on different days of storage, CFU/ml.

The initial TVC increase by Day 3 reflects active LAB growth, followed by a decline due to post-acidification-induced acidity stress (Li et al., 2023). St-Y’s minimal decrease (2.1%) suggests a protective effect, likely from Stevia rebaudiana’s prebiotic enhancement of *Lactobacillus plantarum* survival (Sakr & Massoud, 2021). This aligns with (Plessas et al., 2024), reporting a 2–3% decline in L. plantarum by day 10. TVC above 10^6^–10^7^ CFU/ml meets probiotic standards (Tamime & Robinson, 2007), supporting synbiotic benefits.

### 3.12 Phytochemical analysis

#### 3.12.1 Total Phenolic Content

The total phenolic content (TPC) of yogurt samples (SP-Y, P-Y, St-Y, In-Y) was measured over 10 days of storage at 4°C (Figure 4). Initial TPC (Day 1) averaged 0.67 µg GAE/g (SP-Y), 1.07 µg GAE/g (P-Y), 16.67 µg GAE/g (St-Y), and 0.53 µg GAE/g (In-Y). For St-Y, TPC decreased by 3% to 15.33 µg GAE/g by Day 5 and further to 14.33 µg GAE/g by Day 10 (-14%). SP-Y, P-Y, and In-Y showed minimal variation, with final TPC averaging 0.83, 1.42, and 0.42 µg GAE/g, respectively. Two-way ANOVA confirmed significant effects of storage day (F (4, 40) = 20.12, p = 3.81 × 10^−9^), sample type (F(3,40) = 13929.98, p = 2.10 × 10^−60^), and their interaction (F(12,40) = 14.75, p = 4.14 × 10^−11^).

**Figure 4:**
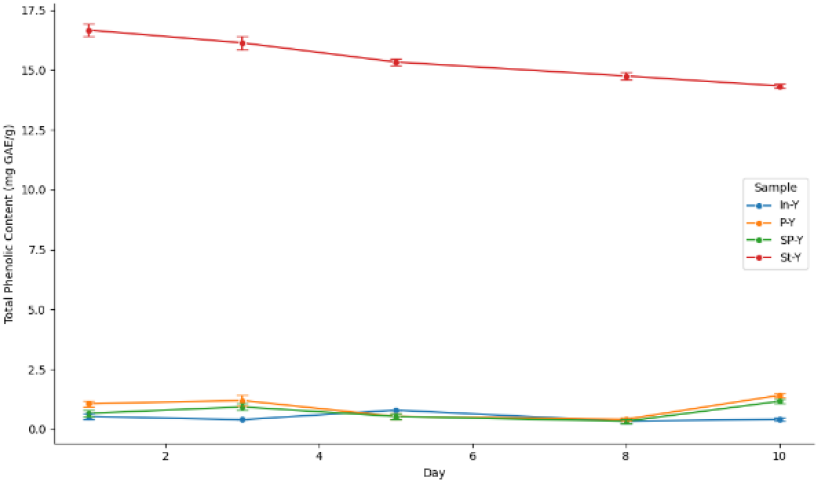
Total phenolic content in yogurt samples during storage.

**Figure 5:**
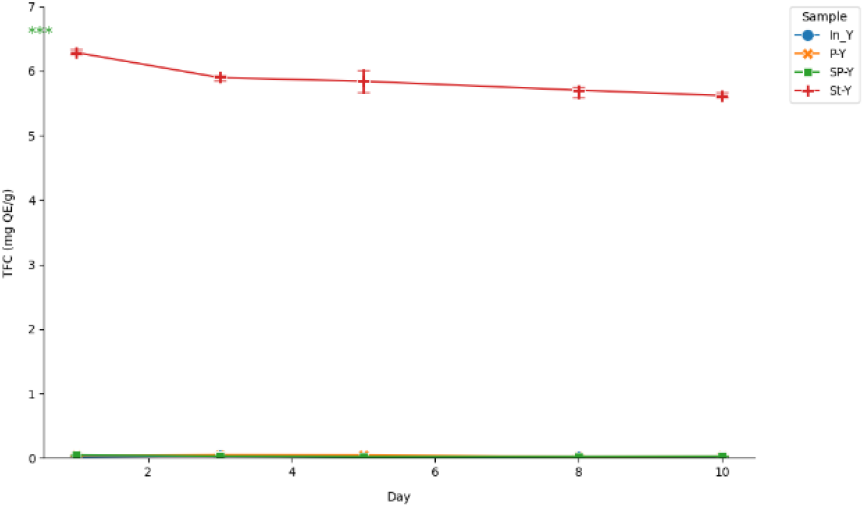
Total Flavonoid Content (TFC) of yogurt samples during storage.

The significant TPC decline in St-Y (-14%) reflects degradation of phenolic compounds from Stevia rebaudiana, known for high phenolic levels (16.67 µg GAE/g) (Grozeva et al., 2015). This aligns with (de Carvalho et al., 2019), who reported stable TPC in stevia-supplemented yogurt after 10 days. The stability of SP-Y, P-Y, and In-Y suggests minimal phenolic contribution, likely due to non-specific Folin-Ciocalteu assay reactions with reducing sugars or proteins (Li et al., 2023; Shori et al., 2018). Phenolic compounds enhance yogurt’s antioxidant potential as free radical scavengers (Khiraoui et al., 2018), with St-Y offering the greatest benefit.

##### 3.12.2 Total Flavonoid Content

The total flavonoid content (TFC) of yogurt samples (SP-Y, P-Y, St-Y, In-Y) was assessed over 10 days of storage at 4°C (Figure 6). Initial TFC (Day 1) averaged 0.048 µg QE/g (SP-Y), 0.040 µg QE/g (P-Y), 6.284 µg QE/g (St-Y), and 0.020 µg QE/g (In-Y). For St-Y, TFC decreased by 7% to 5.895 µg QE/g by Day 5 and further to 5.618 µg QE/g by Day 10 (-11%). SP-Y, P-Y, and In-Y showed negligible changes, with final TFC averaging 0.029, 0.029, and 0.015 µg QE/g, respectively. ANOVA revealed significant differences across samples (p < 0.001), with post-hoc analysis confirming St-Y’s substantially higher TFC compared to SP-Y, P-Y, and In-Y (p < 0.001), while no significant differences were observed among the latter three (p > 0.98).

**Figure 6:**
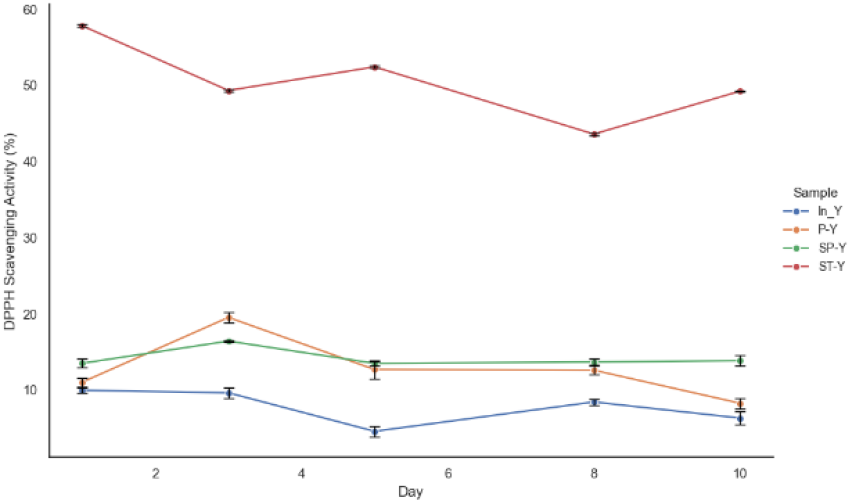
%DPPH in yogurt during storage at 4ºc for 10 days.

The significant TFC decline in St-Y (-11%) reflects flavonoid degradation from *Stevia rebaudiana*, supporting its high initial content (6.284 µg QE/g) (Khiraoui et al., 2018). This aligns with reported ranges of 33.31–50.04 mg RE/gm in stevia extracts (Kim et al., 2011). The negligible TFC in SP-Y, P-Y, and In-Y suggests minimal flavonoid presence, likely due to the aluminum chloride assay’s selectivity for flavones and flavanols, underestimating total flavonoids (Kim et al., 2011).

Flavonoids enhance antioxidant activity, with St-Y offering the greatest potential (Santos-Buelga et al., 2019). Future studies should use HPLC to quantify specific flavonoids and assess storage stability.

##### 3.12.3 DPPH radical scavenging assay

The antioxidant potential of the yogurt samples, measured via the DPPH radical scavenging assay, revealed significant variations across formulations and storage periods. Among the samples, St-Y exhibited the highest initial DPPH radical scavenging activity at 57.84%, followed by SP-Y (13.52%), P-Y (11.05%), and In-Y (9.97%). A two-way ANOVA indicated that sample type significantly influenced DPPH activity (F (3, 40) = 18,640.34, p < 0.0001). Likewise, storage time had a significant effect (F (4, 40) = 153.01, p < 0.0001), and a significant interaction between sample type and storage time was observed (F (12, 40) = 113.52, p < 0.0001).

During refrigerated storage, the antioxidant activity of SP-Y, P-Y, and In-Y declined notably from day 5 and stabilized thereafter. On the 10th day, their activities were reduced to 13.87%, 8.28%, and 6.33%, respectively. In contrast, St-Y showed a slower degradation pattern, with its DPPH activity dropping by approximately 9% by day 5 and further declining to 49.26% on the 10th day, representing a 15% reduction compared to the initial value.

The consistently superior antioxidant performance of St-Y can be attributed to the presence of aqueous stevia extract, which is known for its robust antioxidant properties. This observation is in line with (Sakr & Massoud, 2021), who reported that stevia sweeteners exhibit potent antioxidant activity (∼69.61%) and interact synergistically with probiotics. Additionally, found that the antioxidant activity of aqueous stevia leaf extract ranged from 69.01% to 78.08%, further supporting the current findings. (de Carvalho et al., 2019) also demonstrated that incorporating stevia into yogurt enhances the delivery of health-promoting compounds, thereby enriching the food matrix. Thus, the inclusion of *Stevia rebaudiana* in fermented dairy products such as yogurt presents a promising strategy for developing functional foods with enhanced bioactivity, as also supported by (Khalid et al., 2021) and (Myint et al., 2020).

## CONCLUSION

This study demonstrates the functional potential of synbiotic yogurt enriched with *Lactobacillus plantarum* and aqueous *Stevia rebaudiana* extract. The selected probiotic strain exhibited strong cell surface hydrophobicity, high auto-aggregation, and strain-specific co-aggregation with food-borne pathogens, suggesting potential colonization and pathogen exclusion abilities. However, in vivo validation is essential to confirm survivability and probiotic efficacy under gastrointestinal conditions, which can differ significantly from in vitro environments. Yogurt samples enriched with stevia extract demonstrated enhanced nutritional value, notably higher protein content, suggesting that stevia not only serves as a natural sweetener but also contributes additional

### 3.13 Sensory Evaluation of Products

The sensory attributes of the yogurt samples, including appearance, flavor, taste, mouthfeel, and overall acceptability, were evaluated by a panel of 10 untrained members on Day 1 of storage. Among the formulations, St-Y received the highest ratings for taste (9.2), mouthfeel (9.2), and overall acceptance (8.2), indicating strong consumer preference. In contrast, all samples received comparable scores for flavor (7.4–7.8) and appearance (7.7–7.9), suggesting minimal differentiation in these specific attributes (Figure 7).

**Figure 7:**
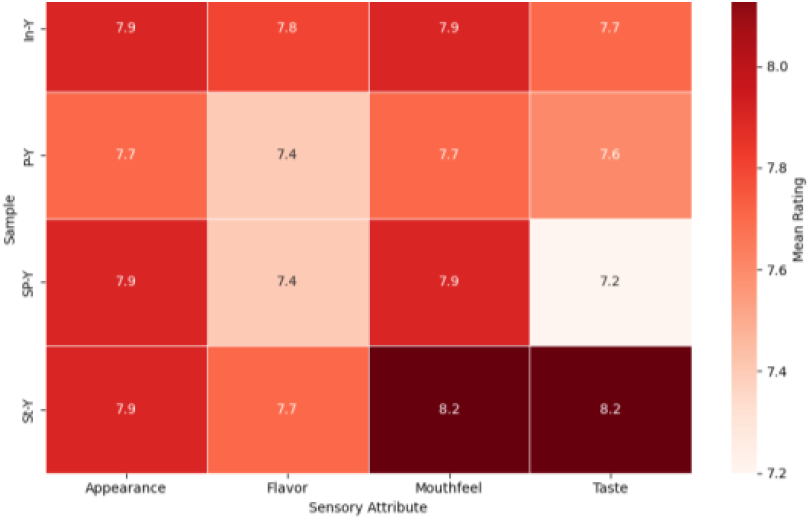
Sensory evaluation of products.

A one-way ANOVA confirmed that overall acceptability significantly differed between samples (F (3,36) = 7.57, p < 0.001). Further Tukey post-hoc tests revealed that St-Y was rated significantly higher than In-Y (p = 0.018), P-Y (p < 0.001), and SP-Y (p = 0.003). No significant differences were observed among the other sample pairs (all p > 0.05). Additionally, St-Y displayed the highest mean acceptability score (8.2 ± 0.48) and greatest panelist variability, whereas P-Y demonstrated complete panelist agreement (7.5 ± 0.0).

The superior sensory performance of St-Y can likely be attributed to the incorporation of *Stevia rebaudiana*, which may enhance both flavor and mouthfeel through its sweetening and bioactive properties. Previous findings by support this interpretation, reporting that stevia extract positively influences the organoleptic and textural qualities of yogurt. These results suggest that the addition of stevia not only contributes to functional benefits such as antioxidant activity but also improves consumer acceptability, making it a promising ingredient for the development of health-oriented dairy products. bioactive components. The synbiotic yogurt (St-Y) exhibited appreciable antioxidant properties, with total phenolic and flavonoid contents of 16.67 µg GAE/g and 6.28 µg QE/g, respectively, and 57.84% DPPH radical scavenging activity on the first day of refrigerated storage. Sensory evaluation indicated that the St-Y variant had superior textural properties, improved palatability, and greater overall acceptability compared to control formulations. Overall, the incorporation of *Stevia rebaudiana* extract and *Lactobacillus plantarum* into yogurt enhanced its nutritional, antioxidant, and sensory properties, suggesting its potential as a functional dairy product. However, further in vivo and clinical studies are needed to validate its efficacy and stability under gastrointestinal conditions.

